# *CHRNA5* links chandelier cells to severity of amyloid pathology in aging and Alzheimer’s Disease

**DOI:** 10.1101/2022.05.03.490491

**Authors:** Jonas Rybnicek, Yuxiao Chen, Milos Millic, Earvin Tio, JoAnne McLaurin, Timothy J. Hohman, Philip L De Jager, Julie A Schneider, Yanling Wang, David A Bennett, Shreejoy Tripathy, Daniel Felsky, Evelyn K Lambe

**Author notes:** **Co-Corresponding Authors email addresses**: Shreejoy Tripathy, Dan Felsky, Evelyn Lambe.

## Abstract

Changes in high-affinity nicotinic acetylcholine receptors are intricately connected to neuropathology in Alzheimer’s Disease (AD). Protective and cognitive-enhancing roles for the nicotinic α5 subunit have been identified, but this gene has not been closely examined in the context of human aging and dementia. Therefore, we investigate the nicotinic α5 gene *CHRNA5* and the impact of relevant single nucleotide polymorphisms (SNPs) in prefrontal cortex from 922 individuals with matched genotypic and *post-mortem* RNA sequencing in the Religious Orders Study and Memory and Aging Project (ROS/MAP). We find that a genotype robustly linked to increased expression of *CHRNA5* (rs1979905A2) predicts significantly reduced cortical β-amyloid load. Intriguingly, co-expression analysis suggests *CHRNA5* has a distinct cellular expression profile compared to other nicotinic receptor genes. Consistent with this prediction, single nucleus RNA sequencing from 22 individuals reveals *CHRNA5* expression is disproportionately-elevated in chandelier neurons, a distinct subtype of inhibitory neuron known for its role in excitatory/inhibitory (E/I) balance. We show that chandelier neurons are enriched in amyloid-binding proteins compared to basket cells, the other major subtype of PVALB-positive interneurons. Consistent with the hypothesis that nicotinic receptors in chandelier cells normally protect against β-amyloid, cell-type proportion analysis from 549 individuals reveals these neurons show amyloid-associated vulnerability only in individuals with impaired function/trafficking of nicotinic α5-containing receptors due to homozygosity of the missense *CHRNA5* SNP (rs16969968A2). Taken together, these findings suggest that *CHRNA5* and its nicotinic α5 subunit exert a neuroprotective role in aging and Alzheimer’s disease centered on chandelier interneurons.

## Introduction

The cholinergic system plays a critical role in the pathology of Alzheimer’s disease (AD) (1), a neurodegenerative disease marked by the accumulation of β-amyloid peptide (β-amyloid) and neurofibrillary tangles of phosphorylated tau in the brain (2). In AD, there are well-characterized disturbances in the excitation/inhibition (E/I) balance in cerebral cortex (3,4) arising from the disruption of inhibitory signalling. The shift toward higher excitation in the cortex is associated with cognitive impairment in AD (5).

The cholinergic system is an important regulator of E/I balance in the prefrontal cortex (PFC) (6,7) and is central to one of the first mechanistic explanations of cognitive deficits in AD; the so-called *cholinergic hypothesis* (8). In AD, there is a decrease of cortical nicotinic acetylcholine receptor binding (9,10), and β-amyloid binding to nicotinic receptors has been postulated as a potential mediator of AD pathology (11,12) possibly via blockade of these receptors (13,14). By contrast, stimulation of neuronal nicotinic receptors has been found to improve survival of primary neuronal culture exposed to β-amyloid (15), and to promote neurogenesis and improve cognition (16,17) in preclinical models. Promoting nicotinic signalling using acetylcholinesterase inhibitors is one of the mainstay AD treatments (18).

High-affinity nicotinic acetylcholine receptors are hetero-pentamer cation channels most commonly composed of α4 and β2 subunits (α4β2) (19–21)(**Fig. 1A**). Deep layer PFC pyramidal cells express nicotinic receptors also containing the auxiliary α5 subunit (22–25). Nicotinic α5 subunits do not contribute to the acetylcholine binding site and cannot form functional receptors on their own (26), requiring the binding sites provided by partner subunits α4 and β2, and forming the α4β2α5 nicotinic receptor. The α5 subunit alters the kinetics of nicotinic receptors (22,23) and increases their permeability to calcium ions (27,28). Importantly, β-amyloid binds less readily to α4β2α5 than α4β2 nicotinic receptors *in vitro* (14), which raises the question of a possible protective role of the α5 subunit in AD pathology.

**Figure 1.**
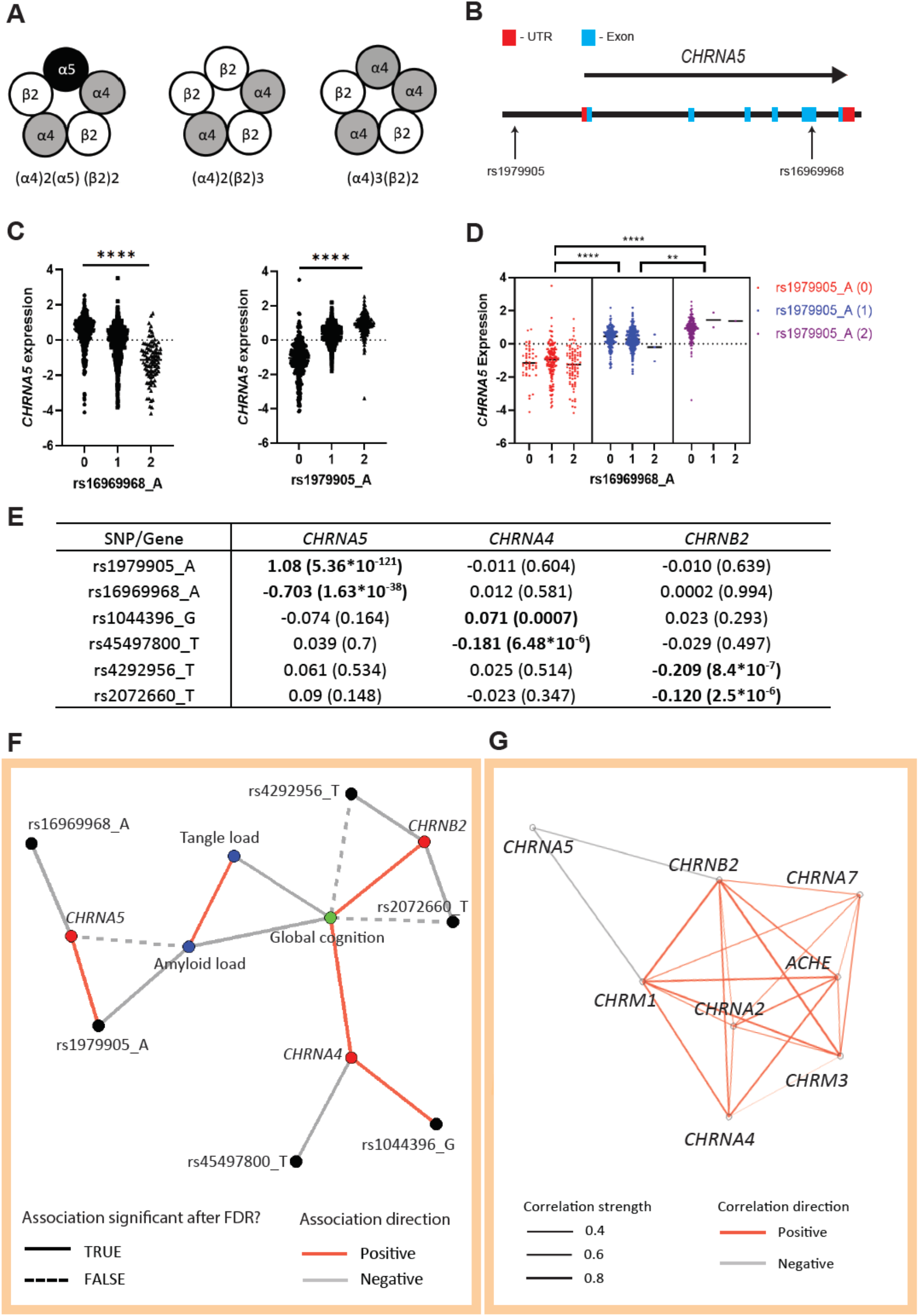
SNPs affecting expression of α4β2* nicotinic receptor subunit genes highlight a link between *CHRNA5* and amyloid pathology. ***A,*** Schematics illustrating different subunit compositions of prefrontal α4β2* nicotinic receptors, with and without the α5 subunit. ***B,*** Localization of the rs16969968 and rs1979905 SNPs in relation to the *CHRNA5* locus*. **C,*** The A allele of the missense SNP rs16969968 (left) in the coding region of *CHRNA5* is associated with lowered *CHRNA5* expression, while the A allele of the rs1979905 SNP (middle) upstream of the *CHRNA5* gene is associated with enhanced *CHRNA5* expression. Data from the DLPFC of ROS/MAP individuals. ***D,*** *CHRNA5* expression appears to be controlled by the zygosity of the rs1979905 A allele (colors) instead of the rs16969968 A allele (x-axis). Data shown as *CHRNA5* expression for each subject with means indicated. ***E,*** eQTL effects of SNPs in nicotinic subunit genes on their respective gene expression. Shown as β-coefficient with significance (p) in brackets. **F,** Network plot depicting the relationships between SNPs (black), gene expression (red), neuropathology (blue), and last global cognition score (green). Solid and dashed lines indicate whether association was significant after correction for FDR or not, respectively. **G,** Network plot depicting the correlations present between the expression of select cholinergic genes in the DLPFC. Colour of lines indicates direction of correlation (negative or positive) while the thickness indicates correlation strength. All correlations shown are significant after adjustment for FDR.

The nicotinic α5 subunit has previously been linked to cognitive performance, with loss or disruption of this subunit impairing performance in attentional tasks in rodents (24,29). In humans, single nucleotide polymorphisms (SNPs) affecting the expression and function/trafficking of the *CHRNA5* gene, which codes for the α5 subunit, have been linked to attentional and cognitive deficits (30,31). These SNPs are also linked to smoking (32), a major AD risk factor (33). However, the role of *CHRNA5* in human aging and AD is unknown.

To address this critical gap, we built a multi-step model of the connections between SNPs affecting the expression and function of *CHRNA5*, and age-related cognitive and neuropathological phenotypes using detailed clinical and post-mortem data from the Religious Order Study and Memory and Aging Project (ROS/MAP) (34). Next, we leveraged single-nucleus RNAseq to determine the cell-type expression pattern of *CHRNA5* in the PFC. We then used a gene ontology analysis of cortical patch-seq data (35) to elucidate the functional makeup of the cell type with the highest *CHRNA5* levels in the PFC, the chandelier cells. Finally, we probed an estimated cell type proportion dataset from the PFC (36) to assess the interaction effect of *CHRNA5* SNPs and Alzheimer’s disease pathology on this *CHRNA5*-enriched cell type. The results of our study suggest a novel role for *CHRNA5* in maintaining the E/I balance in the forebrain and as a potential new target for therapies aiming to promote neuronal survival in AD.

## Methods

### Study cohort

We accessed data from 2004 deceased individuals from the ROS/MAP cohort study (34), of whom 1732 were autopsied. The participants of both studies enrolled without known dementia. ROS enrols elderly nuns, priests, and members of clergy, whereas MAP enrols individuals from community facilities and individual homes. Both studies were approved by an Institutional Review Board of Rush University Medical Centre. Participants gave informed consent for annual clinical evaluation, completed a repository consent allowing their resources to be shared, and signed an Anatomic Gift Act for brain donation at the time of death. Most individuals assessed were female (68%). The average age at study entry was 80.5 ± 0.16 years and the average age at death was 89.2 ± 0.2 years. All data was retrieved from the Synapse AMP-AD Knowledge Portal (Synapse ID: syn2580853).

### Selection of candidate single nucleotide polymorphisms (SNPs)

Nicotinic α5 subunits (encoded by *CHRNA5*) do not contribute to the acetylcholine binding site and cannot form functional receptors on their own (26). In prefrontal cortex, the nicotinic α5 subunits participate in pentameric receptors with two binding sites contributed by partner subunits: α4 (encoded by *CHRNA4*) and β2 (encoded by *CHRNB2*) (19). Together, these subunits form α4β2α5 nicotinic receptors. While the current focus is *CHRNA5*, we also probed the impact of polymorphisms relevant to its receptor partners *CHRNA4* or *CHRNB2* for perspective. The specific polymorphisms were selected based on their reported effects on gene expression, function, or on their clinical associations. For *CHRNA5*, we selected the non-synonymous SNP rs16969968 (minor allele frequency (MAF) = 0.33) in the coding region of *CHRNA5* (**Fig. 1B**) and a cluster of 6 SNPs in linkage disequilibrium (LD) in the regulatory region of *CHRNA5* (37) represented in our study by rs1979905 (MAF = 0.43)(**Fig. 1B**). These SNPs have previously been linked to altered *CHRNA5* expression (37) and altered risk of nicotine addiction (32,38). For *CHRNB2,* the T allele of the rs2072660 polymorphism (MAF = 0.23) has previously been linked to altered risk for nicotine dependence (39), and the T allele of the rs4292956 (MAF = 0.07) with a clinical response to treatment for smoking cessation (40). For *CHRNA4*, the G allele of the rs1044396 polymorphism (MAF = 0.45) has previously been associated with altered performance in attention tasks as well as with the regulation of *CHRNA4* expression (41).

### Genotype data preparation and imputation, quality control, generation of bulk gene expression residuals

Details on the ROS/MAP cohort genotyping and handling of the post-mortem samples have been previously published (42) and are described briefly together with the quality control approaches and generation of the gene expression residuals in **Supplemental Methods**.

### Final clinical diagnosis

We included the final clinical diagnosis as a covariate where relevant in our models. At the time of death, all available clinical data were reviewed by a neurologist with expertise in dementia, and a summary diagnostic opinion was rendered regarding the most likely clinical cognitive diagnosis at the time of death. The methodology of the clinical diagnosis has been previously published (43,44). Summary diagnoses were made blinded to all postmortem data. Case conferences including one or more neurologists and a neuropsychologist were used for consensus on some specific cases. The diagnoses we coded as: 1) NCI: No cognitive impairment (No impaired domains), 2) MCI: Mild cognitive impairment (One impaired domain) and NO other cause of CI, 3) MCI: Mild cognitive impairment (One impaired domain) AND another cause of CI, 4) AD: Alzheimer’s dementia and NO other cause of CI (NINCDS PROB AD), 5) AD: Alzheimer’s dementia AND another cause of CI (NINCDS POSS AD), 6) Other dementia: Other primary cause of dementia.

### Neuropathology and cognitive scores

A detailed description of the neuropathology and cognitive variables in ROS/MAP is included in **Supplemental Methods** and on the RADC Research Resource Sharing Hub.

### Single-nucleus RNA sequencing data processing

The single-nucleus gene counts and metadata available from Cain et al. 2020 (36) on synapse (ID: syn16780177) were converted into a Seurat object (45) in R Studio for further processing. Potential doublets were removed by filtering out cells with over 2500 detected features, and potential dead or dying cells were removed by excluding cells expressing over 5% mitochondrial genes. Cell type annotations were indicated in the data as described in the metadata downloaded from the Cain et al. 2020 Synapse repository (ID: syn16780177). The snRNAseq data was log-normalized and matched with genotype data (n = 22 individuals). *CHRNA5* expression was averaged per cell type per individual and *CHRNA5* levels in different cell types were then compared between cell types by one-way ANOVA with Tuckey’s post-hoc t-test. To prevent bias for rare cell types in calculating average *CHRNA5* expression per cell type per individual, cell types with fewer than 100 individual cells represented in the original data (not aggregated) were removed (the layer 5 FEZF2 ET cell type was excluded from analysis as only 93 cells of this type were present in the genotype-matched snRNAseq dataset). The number of cells per cell-subtype cluster in the genotype-matched snRNAseq dataset can be viewed in the supplementary materials (**Supplemental Table S1**). The identity of chandelier cells was further validated by assessing the expression of SCUBE3, a chandelier cell marker (46), across the cell subtypes.

### Gene ontology of chandelier cell genes

A set of genes which are upregulated in PVALB+ chandelier cells versus PVALB+ non-chandelier cells (basket cells) was previously generated by Bakken and colleagues (35). In short, the authors utilized the patch-seq method (combines electrophysiological patch-clam recording with RNA sequencing and morphological analysis) to characterize the transcriptional differences between cortical PVALB+ chandelier cells, and basket cells across multiple species (human, mouse, and marmoset). In our gene ontology analysis, we filtered the list of differentially expressed genes (DEGs) for those specifically upregulated in human PVALB+ chandelier cells versus human basket cells (remaining DEG n = 222). To determine the ontology of this upregulated gene set we used the Gene Ontology Resource (geneontology.org), querying specifically for molecular function.

### Estimates of relative cell type proportions from bulk DLPFC RNAseq

Estimates of cell-type proportions from bulk DLPFC RNAseq data from 640 ROS/MAP participants was performed and described by Cain et al. 2020 (36). In brief: The authors developed a custom regression-based consensus model, CelMod, to extract cell cluster specific genes from the snRNAseq dataset from 24 ROS/MAP participants, and then used these genes to estimate the proportions of different cell subtypes in the bulk DLPFC RNAseq dataset from 640 ROS/MAP individuals. The proportions were estimated using a large set of marker genes extracted by CelMod for each cell subtype. This method was used to ensure that changes in the expression of a small number of major marker genes would not skew the proportions of any given cell type. The deconvolved cell-type proportion data from the DLPFC was available on request from the research group (Cain et al. 2020 personal communication). We matched the cell type proportion data with genotype, bulk DLPFC RNAseq, and neuropathology (brain levels of β-amyloid and tau) data (final n = 549 individuals). At the time of analysis full cell-type annotations (like that in the snRNAseq data) were only available for the different classes of inhibitory neuron proportions. The estimated proportions of a GABAergic neuron subtype, marked by its co-expression of (among other marker genes) PVALB+/LHX6+/THSD7A+, were determined (Cain et al. 2020 personal communication) to represent chandelier cell proportions in DLPFC. Observed differences in the estimated proportions of this cell subtype represent a difference in the proportion of this cell subtype in the broad cell class (GABAergic neurons). Further information on the cell type proportions is provided in **Supplemental Methods**.

## Statistical approaches

### EQTL and conditional eQTL analyses of candidate SNPs and gene expression in DLPFC

All analyses were performed in R (R version 4.0.2) (R Core Team, 2020) unless otherwise indicated. For eQTL analyses (n = 924), linear regression was used to model batch- and technical covariate-corrected gene expression residuals against SNP allele counts, co-varying for the first 10 genomic principal components (fine population structure), biological sex, age at death, and PMI. Bonferroni correction was applied across tests (6 cis-eQTL tests performed). To assess the effects of the *CHRNA5* haplotypes tagged by rs16969968 and rs1979905 on *CHRNA5* expression (**Fig. 1C, D, E**) we used a nested ANOVA analysis with Sidak’s post-hoc test for multiple comparisons in GraphPad Prism.

### Association analysis between SNPs, gene expression, AD neuropathology, and cognition

To map the multi-omic relationships between candidate SNPs (n_SNPs_= 6), cholinergic receptor gene expression (n_genes_= 3), post-mortem AD-related neuropathologies (n_pathologies_=2), and global cognitive performance at the population level, we fit a series of independent linear models, maximizing sample sizes for each combination of data types, and including appropriate technical and biological covariates. Only individuals with genotype data were included in these models (**Supplemental Fig. S1**). SNPs were modelled using additive allelic dosage coding (i.e., 0,1,2). For the associations of SNPs with β-amyloid and tau pathology (n = 1317), sex, age at death, post-mortem interval (PMI), genotyping batch, the first 10 genomic PCs, final clinical diagnosis and *APOE* genotype were included as covariates. For the association of amyloid with tangle pathology (n = 1317) sex, age at death, PMI, final clinical diagnosis and *APOE* genotype were included as covariates. For the associations of SNPs with global cognitive performance at time of death (n = 1448), sex, age at death, years of education, genotyping batch, and the first 10 genomic PCs were included as covariates. To aid in the interpretation of SNP and gene expression effects in the context of co-occurring neuropathologies and cognitive dysfunction, we also modelled the effects of amyloid and tau measures on the last cognition score in our dataset (n = 1253); sex, age at death, PMI and years of education were used as covariates. For the association of gene expression residuals with neuropathology (n = 914), sex, age at death, PMI, final clinical diagnosis and *APOE* genotype were used as covariates. And for association with global cognition at time of death, we used a linear regression model of *CHRNA4, CHRNA5* or *CHNB2* against the last global cognition score using matched cognition and gene expression data (n = 887). Sex, age at death, PMI, years of education and *APOE* genotype were used as covariates.

In total, we fit 36 models for the network analysis (**Fig. 1F**) All two-sided *p*-values for terms of interest were corrected using the Benjamini and Hochberg false discovery rate (FDR) approach (47) of the p.adjust function in R, with corrected *p*_FDR_<0.05 considered significant.

We used a Chi-square test in GraphPad Prism to study the association between *CHRNA5* SNPs genotype and participant smoking status at baseline.

**Supplemental Fig. S1** describes the specific numbers and exclusions of ROS/MAP in each stage of our analyses.

### Network visualization of multi-omic mapping

To integrate our multi-level association analyses and aid global interpretation, we constructed a network plot summarizing relationships among the different data types analysed (**Fig. 1F**). Network nodes correspond to individual independent and dependent variables, and each edge represents an association derived from one of the linear models described above.

### Cross-correlation of cholinergic gene expression and network construction

To assess the correlation of *CHRNA5* to other major cholinergic genes in the bulk DLPFC dataset, we performed a series of Pearson-correlation analyses on the residualized expression of the different cholinergic genes and used the resulting correlations to construct a correlation network (**Fig. 1G**). Network nodes correspond to the different cholinergic genes and the edges represent the size of the correlation coefficient (r). Only correlations significant after adjustment for false discovery rate (56 comparisons) are shown in the network graph.

### Analysis of single-nucleus RNAseq data from DLPFC

The single-nucleus RNAseq dataset published by Cain et al. 2020 contained cells clustered into 37 subtypes. As we were interested in genotypic effects on *CHRNA5* expression in neurons specifically, to narrow the scope of the analysis, non-neuronal cell subsets (microglia : Micro.1 – Micro.6, oligodendrocyte:Olig.1 – Olig.4, endothelial cells: Endo.1 – Endo.4, astrocyte cells: Astr.1 – Astr.5) were grouped into cell-subtype categories (Microglia, Oligodendrocytes, Endothelial cells, Astrocytes) while neuronal cells were kept in their subtypes (except the L5 RORB IT subtype which was formed by grouping the Exc.Exc.RORB_L5_IT_1 and Exc.Exc.RORB_L5_IT_2 subtypes) as defined in Cain et al. 2020. Using the aggregate function in R, the log-normalized DLPFC snRNAseq *CHRNA5* expression (n = 24) was averaged per cell type (subtype) per individual (using the unique participant "projid" IDs) as follows: [aggregate(*CHRNA5* ∼ projid + subtype, data, mean)].

To assess the expression of *CHRNA5* across the different cell types, we used a one-way ANOVA with Tukey’s post-hoc t-test on the *CHRNA5* expression previously averaged per cell type per individual.

To assess the association between the rs1979905 genotype and *CHRNA5* expression in the DLPFC snRNAseq data, we matched the *CHRNA5* expression data averaged per cell type per individual with genotype data (final = 22). We then used general linear models covaried for sex, age at death, PMI and genotyping batch to assess the association between the rs1979905 genotype and *CHRNA5* expression in the different cell types.

### Interaction modelling of neuropathology, CHRNA5 genotype, and bulk-estimated cell type proportions

To assess the association between amyloid and tau neuropathology, and chandelier cell proportions, we used a linear regression model of these variables covaried for sex, age at death and PMI. A linear regression model with the same covariates was also used to assess the association between *CHRNA5* expression and chandelier cell proportions. To test the association between rs1979905 and rs16969968 genotype with chandelier cell proportions, we used a linear regression model covaried for sex, age at death, post-mortem interval, *APOE* genotype and final clinical diagnosis. Finally, we used a linear interaction model to test the association between genotype-neuropathology interaction on the proportion of chandelier cells, covaried for sex, age at death, post-mortem interval, *APOE* genotype, final clinical diagnosis and the first 10 genomic PCs: [Cell type proportion ∼ Genotype * Neuropathology + sex + age at death + PMI + *APOE* genotype + diagnosis + PC1 + … + PC10].

## Results

### Expression of α4β2α5 receptor component genes is affected by single nucleotide polymorphisms

To identify effects in aging and dementia of gene variants previously shown in younger adults to influence *CHRNA5* expression (37) and α5 coding (27,48), we examined brain expression quantitative trait loci (eQTL). The variants were in weak-moderate linkage disequilibrium in our European ancestry sample (r^2^=0.34), in agreement with previous work (37). We also found that dosage of the A allele of the missense SNP rs16969968 (minor allele frequency (MAF) = 0.33) in the coding region of *CHRNA5* (**Fig 1B**) was associated with lower *CHRNA5* expression (t = - 13.61 *p* = 1.63*10^-38^), consistent with existing data (49). A different SNP haplotype in the regulatory region upstream of *CHRNA5* (**Fig 1B**), denoted here by the A allele of the tag SNP rs1979905 (MAF = 0.43), was associated with higher *CHRNA5* expression (t= 27.47, *p* = 5.36*10^-121^) (**Fig. 1C**). Furthermore, analyses of the coding-SNP rs16969968 and the regulatory-SNP rs1979905 together (**Fig. 1D**) showed that *CHRNA5* expression is predominantly regulated by the regulatory-SNP rs1979905 rather than the coding-SNP, rs16969968, as all rs1979905 A allele non-carriers showed similar levels of *CHRNA5* mRNA regardless of the rs16969968 A allele (Nested one-way ANOVA: F(2,6) = 229.6; Šidák’s post-hoc test for multiple comparisons: rs1979905 A1 vs. A0 *p* = 8*10^-6^, A2 vs. A1 *p* = 0.004, A2 vs A0 *p* = 4*10^-6^), suggesting that the effect of rs1979905 A on *CHRNA5* expression is independent of the rs16969968 SNP genotype. This was also demonstrated using a conditional eQTL model, where the effect of rs16969968 A allele on *CHRNA5* expression was lost when co-varying for rs1979905 A allele (rs16969968A: t = -1.027, p = 0.304; rs1979905A: t = 21.927, *p* = 8.194*10^-86^). No association between *CHRNA5* expression and disease state was detected (**Supplemental Fig. S2**). No trans-eQTL effects were detected (**Fig. 1E**) between either of these SNPs and the expression of required partner nicotinic subunit genes, *CHRNA4* and *CHRNB2*.

To assess the α4 and β2 nicotinic subunits required for the formation of α5-containing α4β2α5 receptors, we extended our eQTL analyses to SNPs in *CHRNA4* and *CHRNB2*, focusing on those associated with altered gene expression or clinical effects(40,41,50). Without exception, eQTL SNP effects for these genes were weaker than those of rs16969968 and rs1979905 for *CHRNA5*. The T allele of *CHRNB2* intronic variant rs2072660 (MAF = 0.23) was associated with lower *CHRNB2* expression (t = -4.738, *p* = 2.5*10^-6^), and a similar association with *CHRNB2* expression was seen with the T allele of the *CHRNB2* non-coding variant rs4292956 (MAF = 0.07) (t = -4.961, *p* = 8.4*10^-7^) (**Fig. 1E**). For *CHRNA4,* the G allele of missense variant rs1044396 (MAF = 0.45) in the coding region of *CHRNA4* was associated with higher *CHRNA4* expression (t = 3.416, *p* = 0.0007). We also used the Gene Query function of the xQTLServe online tool (DLPFC of 534 ROS/MAP participants) to identify the T allele of the intronic variant rs45497800 as associated with decreased *CHRNA4* expression (t = -6.92, *p* = 1.32 * 10^-11^). We then replicated this association in our larger cohort of 924 ROS/MAP individuals (MAF = 0.07) (t = -4.537, *p* = 6.48*10^-6^) (**Fig. 1E**). No associations were found between these SNPs and the expression of other high-affinity nicotinic receptor subunit genes (**Fig. 1E**).

### *CHRNA5* polymorphisms are not associated with smoking status in this largely non-smoking population

The percentage of participants identified as never, previous, and current smokers (**Supplemental Methods**) was 68.2, 29.4, and 2.4 respectively. To investigate previously-reported (30,51) associations between genotype for the *CHRNA5* SNPs and smoking status at baseline (current/former/never smoked), we used a Chi-squared test and found no relationship (rs1979905_A : χ2(4) = 1.575, p = 0.813; rs16969968_A: χ2(4) = 1.317, p = 0.858) in this largely non-smoking population. Smoking status was not used in further analysis, unless specifically indicated.

### A *CHRNA5* polymorphism is negatively associated with brain β-amyloid levels in ROS/MAP

To address interrelationships among nicotinic subunit expression, nicotinic SNPs, and neuropathological and cognitive phenotypes, we used clinical and post-mortem data in a multi-step model, the inclusion and exclusion criteria for this model can be found in **Supplemental Fig. S1**. As illustrated in **Fig. 1F**, both β-amyloid and tau pathology were negatively associated with global cognitive performance proximal to death (tau: t = -18.87 *p* = 5.77 *10^-70^; amyloid: t = -7.88, *p* = -6.99*10^-15^) and positively associated with each other (t = 12.59, *p* = 2.14*10^-34^). Of the SNPs examined, only the SNP increasing *CHRNA5* expression had a significant association with AD neuropathology, with the A allele of the regulatory-SNP rs1979905 significantly negatively associated with β-amyloid load after false discovery rate (FDR) correction for multiple comparisons (t = -2.81, *p*_FDR_ = 0.015, corrected for 36 models per (47)). Intriguingly, expression of the *CHRNA5* gene itself showed a direct negative association with β-amyloid load prior to FDR correction (t = -2.23, *p* = 0.026), whereas the expression of the major nicotinic subunit genes *CHRNA4* and *CHRNB2* showed significant positive associations with the last global cognition score, which remained significant after FDR correction (*CHRNA4*: t = 2.98, *p_FDR_* = 0.013; *CHRNB2*: t = 3.43, *p_FDR_* = 0.003). Conversely, the rs2072660 T allele, associated with lower *CHRNB2* expression, was negatively associated with the last global cognition score prior to FDR correction (t = -1.99 *p* = 0.047). A similar negative association with the last global cognition score (t = -2.3, *p* = 0.022) was found for the T allele of the *CHRNB2* SNP rs4292956.

We extended our assessment of the relationship between *CHRNA5* SNPs and amyloid pathology into the Alzheimer’s Disease Neuroimaging Initiative (ADNI) dataset, which uses different measurements of amyloid levels using brain positron emission topography (PET) or spinal tap sampling of cerebrospinal fluid (**Supplemental Methods**). Preliminary ADNI examination (**Supplemental Methods**) did not detect an association between the *CHRNA5* SNPs rs16969968 and rs1979905 and imaging or spinal tap measures of beta-amyloid (data not shown).

### *CHRNA5* expression does not tightly correlate with other components of the cholinergic system

To further investigate the interrelationships among *CHRNA5* and the major nicotinic subunits as well as other components of the cholinergic system, we performed a separate series of gene expression correlation analyses. Most of the major components of the cholinergic system which were detected in bulk DLPFC data of the ROS/MAP individuals (*CHRNA2*, *CHRNA4*, *CHRNA7*, *CHRNB2*, *CHRM3*, *CHRM1* and *ACHE*) showed significant positive correlation with each other (**Fig. 1G**). By contrast, *CHRNA5* stood out as showing no positive correlation with any of the other major cholinergic genes and only weak negative correlations with the expression of *CHRNB2* and *CHRM1* (**Table 1**). Considering that the expression of *CHRNA5*, *CHRNB2,* and *CHRNA4* are required for the assembly of the high-affinity α4β2α5 nicotinic receptor, it was surprising to see that *CHRNA5* expression was not positively correlated with either *CHRNA4* or *CHRNB2* (**Fig. 1G** and **Table 1**). Therefore, we next investigated whether this lack of correlation may arise from differences in the cell-type specific expression of *CHRNA5* compared to the major nicotinic receptor subunits, *CHRNA4* and *CHRNB2,* which are more broadly expressed.

**Table 1.**
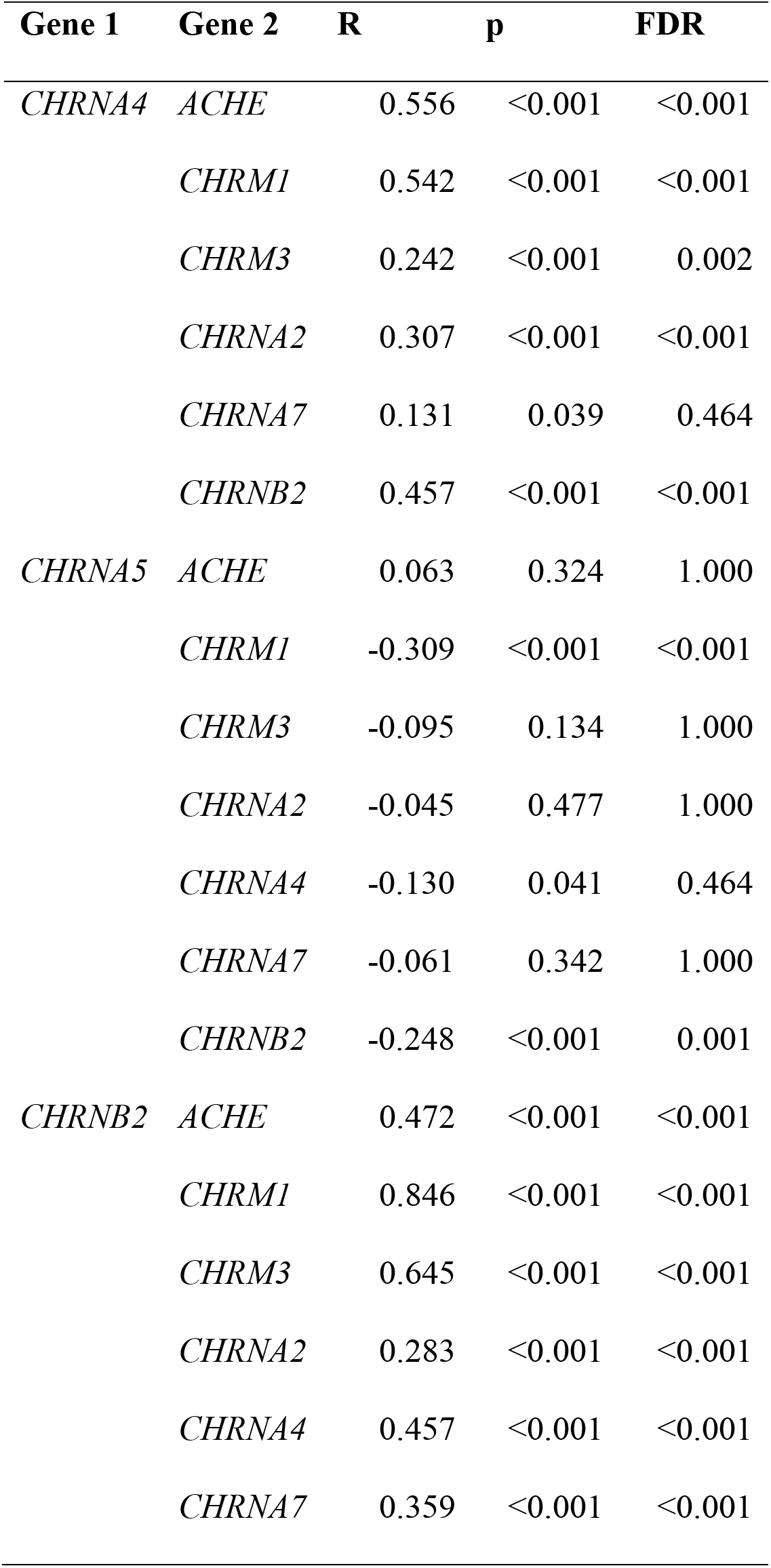
Expression correlation of selected cholinergic genes. Results of Pearson’s correlation analysis of the expression of selected cholinergic genes using the bulk tissue RNAseq data from the DLPFC of ROS/MAP subjects.

### *CHRNA5* shows stronger expression in chandelier interneurons than most other cell classes

To assess the cell-type specificity of *CHRNA5* expression in the ROS/MAP cohort, we calculated the average *CHRNA5* expression per cell type per individual using the genotype-matched single-nucleus RNAseq data available from the DLPFC in a subset of 22 ROS/MAP participants (36)(2 individuals lacked genotyping data). In this small dataset, *CHRNA5* was expressed at a low level across a number of different excitatory, inhibitory, and nonneuronal cell types (**Fig. 2A**), with significantly higher expression in inhibitory PVALB+ chandelier cells (as identified by Cain et al. 2020). Chandelier cells had significantly-higher expression of *CHRNA5* compared to most other cell types (One-way ANOVA: F(21,419) = 3.439, *p* = 6.1*10^-7^; Tukey’s post-hoc t-test Chandelier cells vs. 18 out of 19 other cell types: *p* < 0.05) (**Fig. 2A**). By contrast, chandelier cell expression of *CHRNA4* and *CHRNB2* were at a similar level in chandelier cells to their expression levels in many other cell types (**Fig. 2B**). To confirm the identity of the chandelier as defined by Cain et al 2020 (36), we assessed the expression of *SCUBE3*, a specific marker of chandelier cells (46), across the different cell type found in the ROS/MAP snRNAseq dataset, and found the highest expression of this gene in the chandelier cells (**Supplemental Fig. S3**).

**Figure 2.**
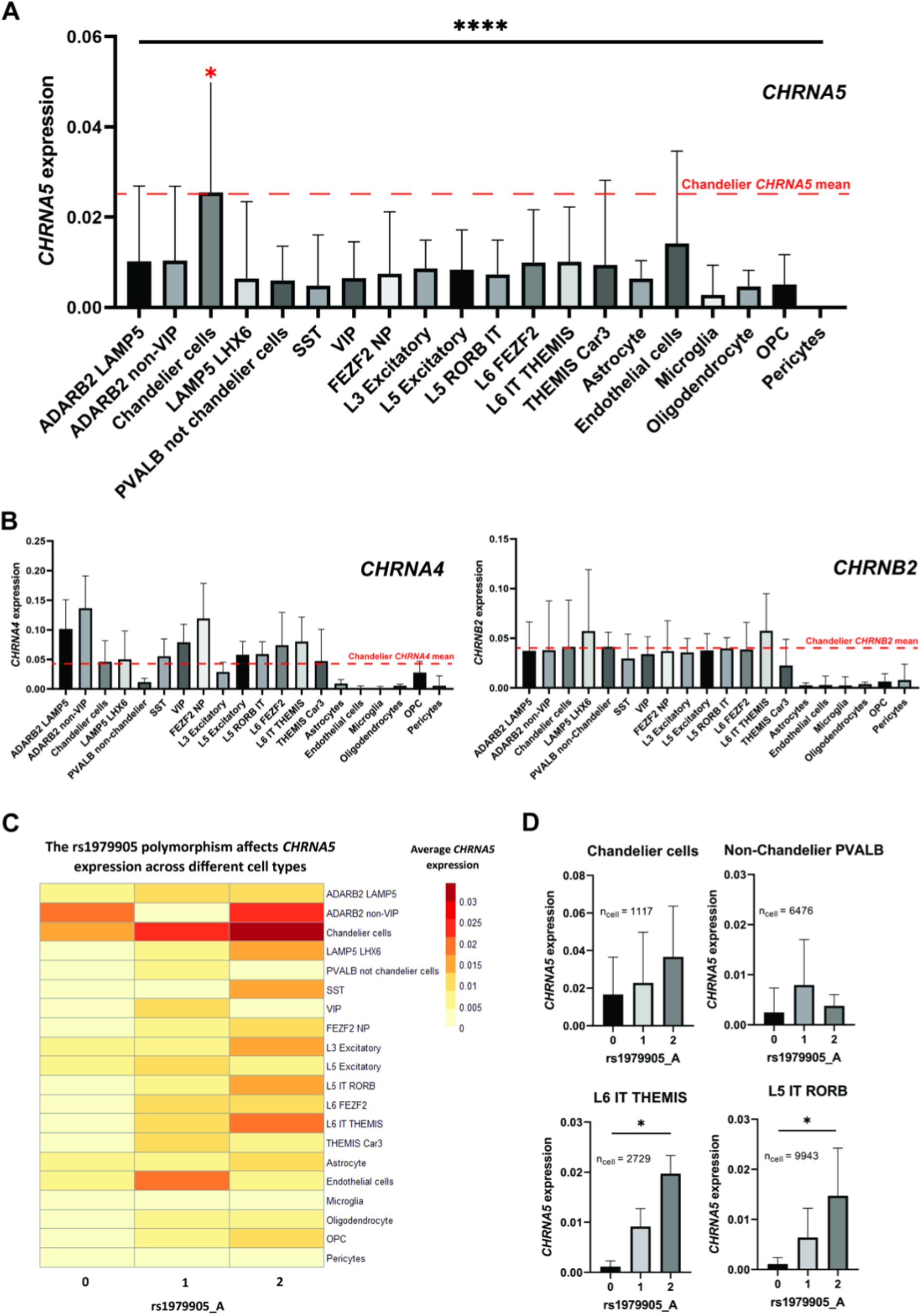
*CHRNA5* expression is elevated in chandelier cells and is affected by genotype for the rs1979905 A allele. ***A,*** Expression of *CHRNA5* averaged per cell type per individual, original gene count values were normalized for each cell by total expression. F-test significance of ANOVA shown on graph, with red asterisk denoting post-hoc tests demonstrating *CHRNA5* expression is stronger in chandelier cells compared to all but one other cell type. Mean expression of *CHRNA5* in chandelier cells displayed (red line). ***B,*** Expression of *CHRNA4* (left) and *CHRNB2* (right) across different cell types in the ROS/MAP DLPFC snRNAseq dataset. Mean expression of *CHRNA4* or *CHRNB2* in chandelier cells displayed (red line). Data shown as mean + SEM of the data averaged per cell-type per individual. ***C,*** Expression of *CHRNA5* across cell types in the PFC in individuals split by genotype for the rs1979905 A allele, expression averaged per cell type per individual. Number of individuals per rs1979905 A allele genotype: 0, n = 5; 1, n = 13; 2, n = 4. ***D,*** Effect of genotype for rs1979905 A allele on expression of *CHRNA5* in selected cell types, data shown as *CHRNA5* expression averaged per cell type per individual. A pattern of increasing *CHRNA5* expression with increasing rs1979905 A zygosity is present in some cell types. Significance shown for linear regression models for L5 and L6 excitatory neurons. Displayed as mean + SEM. Number of cells per subtype indicated (n_cell_).

To confirm the novel finding that chandelier cells show stronger *CHRNA5* expression compared to other classes of neurons, we probed publicly available cell-type specific gene expression databases of human brain tissue. Using the Allen Institute SMART-seq single-cell transcriptomics data from multiple cortical areas https://celltypes.brain-map.org/rnaseq/human_ctx_smart-seq we found *CHRNA5* expression to be highest in a PVALB+/SCUBE3+ inhibitory cell type (0.06 trimmed mean *CHRNA5* expression) likely representing chandelier cells (46), and in a co-clustering PVALB+/MFI+ cell type (0.06 trimmed mean *CHRNA5* expression). Highest expression of *CHRNA5* in chandelier cells compared to all other cell types is also replicated in the Seattle Alzheimer’s Disease Brain Cell Atlas (https://knowledge.brain-map.org/data/5IU4U8BP711TR6KZ843/2CD0HDC5PS6A58T0P6E/compare?cellType=WholeTaxonomy&geneOption=CHRNA5&metadata=CognitiveStatus&comparison=dotplot.) In the Human Protein Atlas database (brain single cell tissue) https://www.proteinatlas.org/ENSG00000169684-CHRNA5/single+cell+type/brain, *CHRNA5* showed highest expression in a PVALB+ inhibitory cell type (c-41) which also showed highest expression of *SCUBE3* (Inhibitory neurons c-41, 15.1 normalized *CHRNA5* transcripts per million), likely also representing chandelier cells.

To investigate the cell type-specificity of the *CHRNA5* eQTL effects of the regulatory-SNP rs1979905 in the single nucleus data from ROS/MAP, we stratified the averaged *CHRNA5* expression by genotype for the rs1979905 A allele. We found that higher allelic dosage of the rs1979905 A allele was associated with greater *CHRNA5* expression (**Fig. 2C**, **Supplemental Table S2**), and that this pattern was most pronounced in subtypes of layer 5 (L5 RORB IT: t = 2.460, *p* = 0.0249) and layer 6 (L6 IT THEMIS: t = 2.402, *p* = 0.028) excitatory neurons (**Fig. 2D, Supplemental Table S2**). In the stronger *CHRNA5*-expressing PVALB+ chandelier cells, however, the association did not reach significance. Other neuronal cell types appeared to diverge completely from the typical eQTL pattern of rs1979905 (**Fig. 2C,D, Supplemental Table S2**) In non-neuronal cells, this eQTL pattern of 1979905 was limited to oligodendrocytes and oligodendrocyte precursor cells (**Supplemental Table S2**). This analysis suggests that only a subset of cell-types contribute to the stepwise expression pattern observed in the prefrontal cortex by rs1979905 genotype.

Cell type specific data from ROS/MAP and the Allen database supports the hypothesis that *CHRNA5* possesses a distinctive expression pattern with enrichment in chandelier cell interneurons, compared to its more abundant and widely-expressed subunit partners.

### Chandelier cells are significantly enriched for genes interacting with β-amyloid

To investigate the potential contribution of chandelier cells to β-amyloid processing, especially one that might be driven by nicotinic α5-containing receptors, we assessed the molecular function of genes which are significantly upregulated in chandelier cells compared to PVALB+ non-chandelier cells (basket cells). For this analysis we took advantage of a list of 222 such genes previously generated by Bakken and colleagues (35), who investigated the cellular identities of chandelier neurons in the human cortex. Gene ontology analysis revealed this gene list to be significant enriched for genes defined as “amyloid-beta binding” (fold enrichment = 7.79, *p*_FDR_ = 0.01) (**Fig. 3A**), including *SORL1,* an endocytic receptor that directs the amyloid precursor protein away from the amyloidogenic pathway (52–54) and *EPHA4,* a receptor tyrosine kinase involved in amyloid regulation (55). Since α5-containing nicotinic receptors are highly permeable to calcium ions (27,28), we note that the gene ontology analysis also showed chandelier neurons to be significant enriched for genes with a “calcium ion binding” molecular function (fold enrichment = 3.77, *p*_FDR_ = 4.12*10^-6^) (**Fig. 3B)**, including genes potentially protective against amyloid pathology such as *MME and SPOCK1*. Two genes are at the intersection of these functional pathways in chandelier cells, *PRNP and CLSTN2*. The former has been implicated in nicotinic receptor-mediated regulation of amyloid (56–59) and the latter has been implicated in cortical GABAergic neurotransmission (60) These findings underscore the importance of investigating the relationship between the functional status of nicotinic α5 subunits and chandelier neuron vulnerability to AD neuropathology.

**Figure 3.**
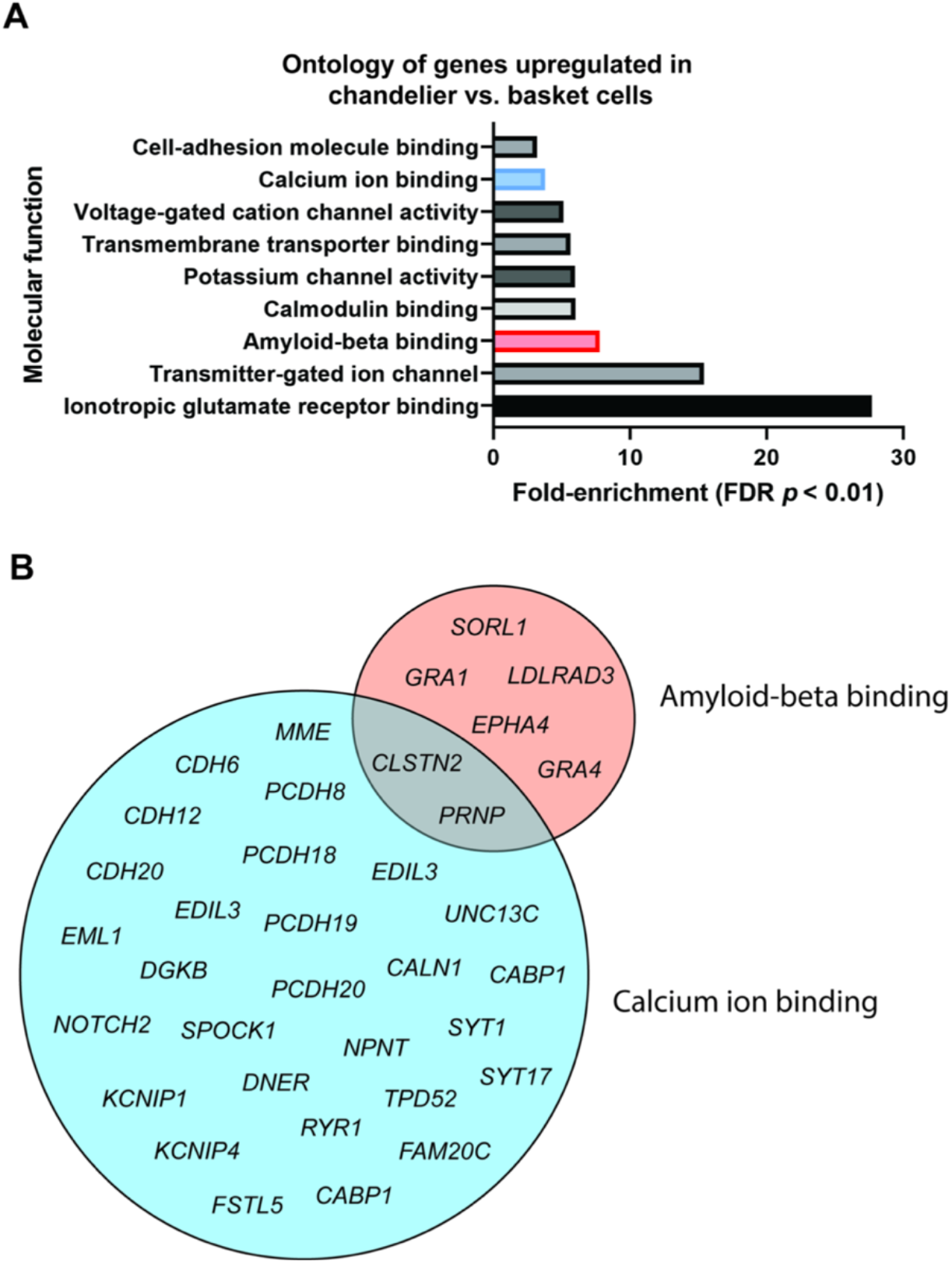
Chandelier cells are significantly enriched for genes interacting with amyloid. ***A,*** Ontology (molecular function) of gene set upregulated in cortical PVALB+ chandelier cells versus PVALB+ non-chandelier cells (basket cells). Only molecular functions with significant fold enrichment after FDR are displayed. “Calcium ion-binding” and “amyloid-beta binding” functions are highlighted. ***B,*** Venn diagram displaying genes from ***A*** with either a “calcium ion-binding” or a “amyloid-beta binding” molecular function, and their overlap.

### A genotype-specific reduction in proportion of chandelier cells with increasing brain β-amyloid levels

To determine whether impaired function/trafficking α5-containing nicotinic receptors might promote chandelier neuron vulnerability to neurodegeneration, we examined the interaction of rs16969968 genotype and AD neuropathology on estimated proportions of chandelier neurons in the bulk RNAseq dataset. This investigation was based on cell type proportion estimates for chandelier cells and several other interneuron subclasses, from sets of single-cell-informed marker genes. The cell type estimates were derived from the bulk DLPFC RNAseq data of a subset of 640 ROS/MAP participants. Overall, β-amyloid levels were negatively associated with the proportion of chandelier cells (t = -4, *p* = 7.26*10^-5^). However, the missense rs16969968 A allele homozygotes showed significantly lower chandelier cell proportions with increasing β-amyloid load, compared to rs16969968 A allele non-carriers (interaction term t = -2.842, *p* = 0.005) (**Fig. 4A,B,C**). In a secondary analysis, there is a suggestion of an opposite relationship of rs1979905 A allele and β-amyloid levels with chandelier cells but this does not reach statistical significance (interaction term t = 1.81, *p* = 0.071) (**Fig. 4D,E,F**). The observed relationships between chandelier cell proportions, *CHRNA5* genotypes and amyloid were not altered by the inclusion of smoking status as a covariate in the analysis. Chandelier cell proportions did not correlate with tau pathology (t = -0.4, *p* = 0.687), nor was there an interaction of *CHRNA5* genotype and tau pathology with chandelier cell proportions (data not shown). To assess whether the interaction between β-amyloid levels and *CHRNA5* SNPs was driven by the effects of these SNPs on *CHRNA5* expression, we assessed the effect of the interaction of *CHRNA5* levels and β-amyloid load on chandelier cell proportion but found no significant effect (interaction term, *t* = 0.5, *p* = 0.619). This suggests that the genotype-specific association between amyloid load and chandelier cell proportion is more likely driven by changes in nicotinic α5 protein structure and/or trafficking (61) as a consequence of having two copies of the missense SNP in *CHRNA5*, rather than by the altered *CHRNA5* expression associated with the rs1979905 SNP genotype.

**Figure 4.**
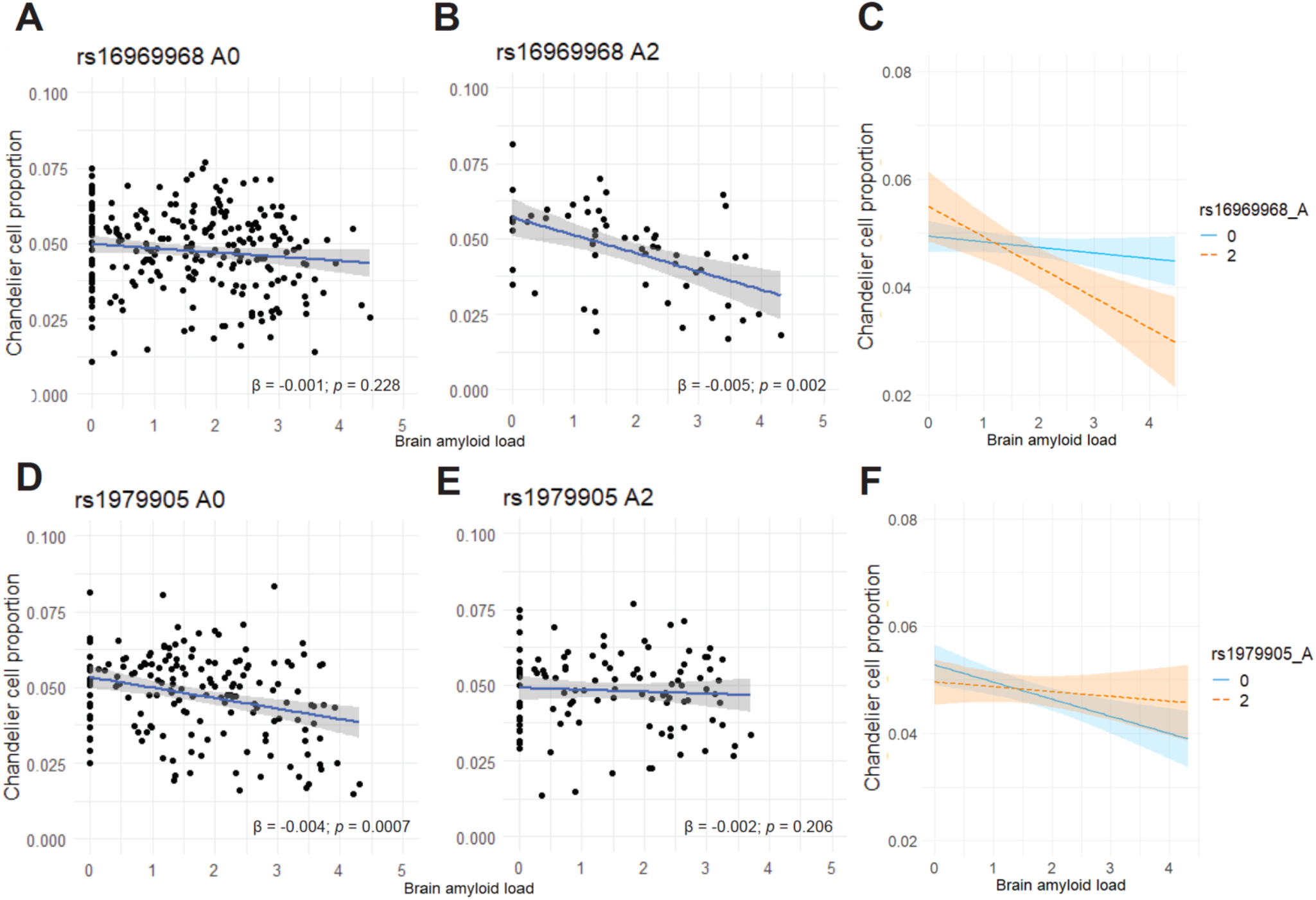
Association of chandelier cell proportions with β-amyloid load is dependent on the rs16969968 A allele genotype. Cell type proportion data for interneuron populations is available for almost a third of the deceased ROS/MAP subjects, allowing the assessment of the interaction among chandelier cell proportion, brain amyloid load, and the *CHRNA5* SNP haplotype. ***A,B*** Stratifying by rs16969968 A allele reveals a significant interaction effect between rs16969968 A allele and amyloid load on chandelier cell proportions with rs16969968 A allele homozygotes showing a more negative association between brain amyloid load and chandelier cell proportions compared to rs16969968 A allele non-carriers (interaction term t = -2.842, *p* = 0.005). Scatter plots show 95% confidence intervals of linear model predictions, β-coefficients and *p* values of individual linear regression models are displayed. ***C***, Overlay of the linear models from A,B, showing 95% confidence intervals. ***D,E,*** Stratifying by rs1979905 A allele shows a suggestion of an opposite interaction between rs1979905 A allele and amyloid load on chandelier cell proportions (interaction term t = 1.81, *p* = 0.071). Scatter plots show 95% confidence intervals of linear model predictions, β-coefficients and *p* values of individual linear regression models are displayed. ***F***, Overlay of the linear models from D,E, showing 95% confidence intervals.

The schematic in **Fig. 5** illustrates a working model of the impact of the rs16969968 A allele homozygosity for chandelier cell vulnerability, as well as example mechanisms enriched in chandelier cells and known to alter β-amyloid processing. **Supplemental Table S3** summarizes the impacts of the regulatory and functional SNPs of *CHRNA5*.

**Figure 5.**
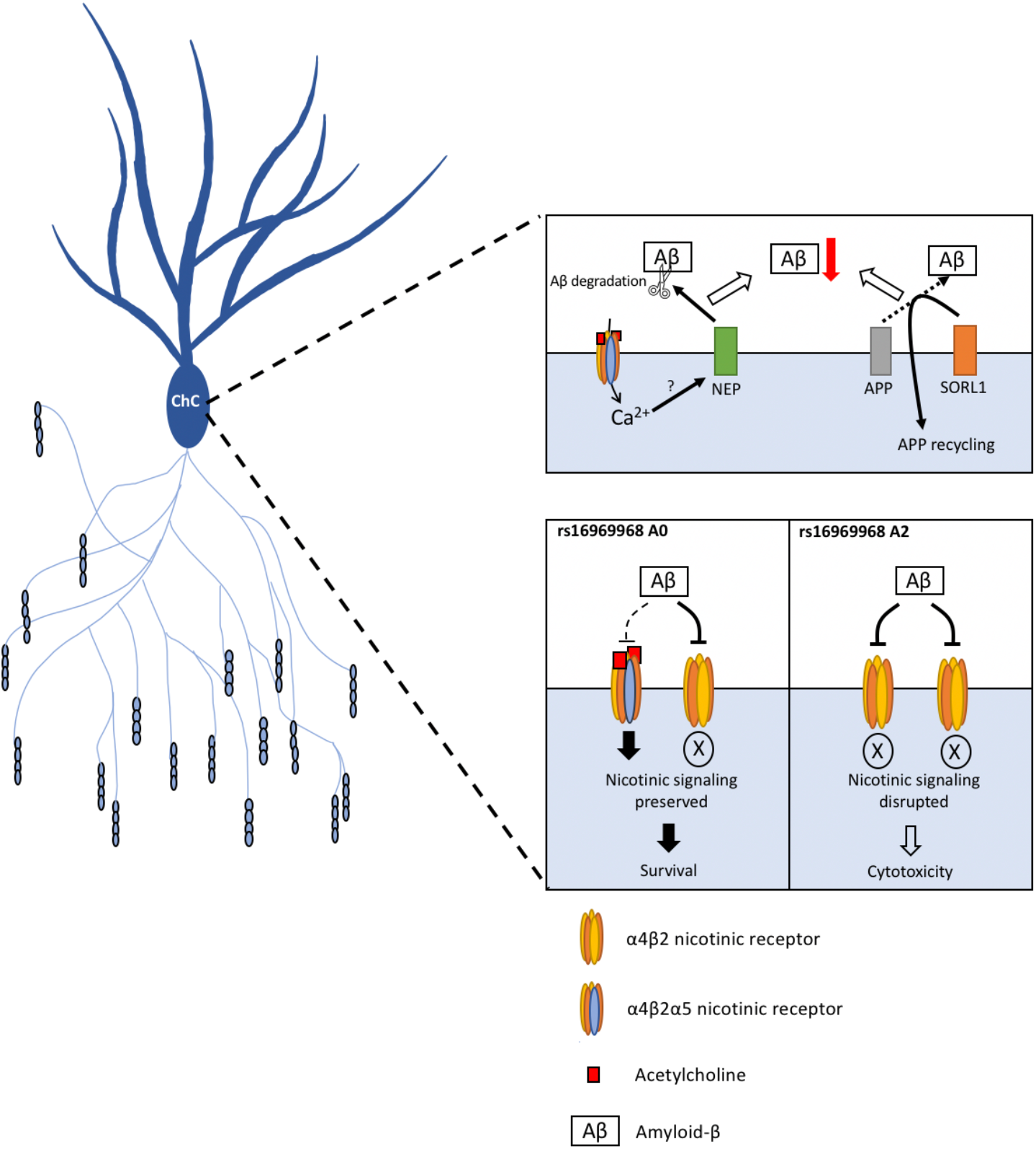
A working model of the potential role of chandelier cells in β-amyloid processing, and of the impact of *CHRNA5* genotype on chandelier cell (ChC) resilience and vulnerability. Top: Chandelier cells are significantly enriched for multiple genes involved in β-amyloid processing and degradation including for example neprilysin (NEP), a potentially *CHRNA5*-regulated degrader of β-amyloid, and SORL1, a vital component of the APP-recycling pathway. Bottom: In coding-SNP rs16969968 non-carriers, the α4β2α5 nicotinic receptor is resistant to inhibition by β-amyloid, preserving nicotinic signalling even at high β-amyloid levels that inhibit the α4β2 receptors. In coding-SNP rs16969968 homozygous individuals, the disruption of the α5 subunit may reduce its representation in the receptors or block its protective function against β-amyloid, leading to disrupted nicotinic signalling at higher β-amyloid levels, possibly triggering a cytotoxic response in the chandelier cells. Chandelier cell structure is based on images from Tai et al. 2014 (91).

## Discussion

We examined human prefrontal cortical nicotinic α5 subunit expression for the first time in aging and AD-related neuropathology. We took advantage of SNPs affecting *CHRNA5* expression and function/trafficking. The aging prefrontal cortex demonstrates strong eQTL effects of the common regulatory-SNP rs1979905 and its A allele is associated with lower levels of brain β-amyloid. Single-nucleus RNAseq data revealed that chandelier cells have the greatest abundance of *CHRNA5* expression in human prefrontal cortex. These neurons are significantly enriched in amyloid-binding proteins, including some that may be activated via nicotinic receptors. We find the common coding-SNP rs16969968 that disrupts nicotinic α5 subunits renders this population of chandelier interneurons population vulnerable to β-amyloid levels. Our findings are summarized in the working model in **Fig 5**, as well as **Supplemental Table S2**. Taken together, they raise the possibility of a cell-type specific neuroprotective role for C*HRNA5* to reduce β-amyloid levels and toxicity.

Neither rs1979905 nor rs16969968 has been significantly associated with Alzheimer’s disease risk in the largest and most recent genome wide association study (GWAS) on late-onset AD (62) (**Supplemental Table S4**). This highlights the value of targeted phenotype-oriented studies like our own, since they allow for the probing of more detailed associations than permitted in GWAS.

### Inhibitory signalling is disrupted in Alzheimer’s disease

The disruption of E/I balance in the cortex is a hallmark of AD pathology and is associated with cognitive AD symptoms (3,4). Previous studies have shown disruption of cortical inhibitory signalling in AD stems from the alteration of the activity of inhibitory neurons and inhibitory cell loss (63–65). However, the susceptibility of inhibitory neurons to AD pathology is not uniform. While studies have shown significant drops in somatostatin-positive interneurons in the cortex of AD patients (66,67), the numbers of parvalbumin-positive cortical interneurons, including the PVALB+ chandelier cells, appear comparatively more resilient to AD pathology (67,68). Our study expands on these findings, demonstrating that the preservation of chandelier cells in AD pathology depends on the genotype of SNP in a nicotinic receptor subunit. The rs16969968 SNP is a missense mutation found to functionally alter the α5-containing nicotinic receptors in cell systems and *in vivo*, through altered channel biosynthesis, trafficking, properties and modulation (27,48,61,69,70). Preclinical opto-physiological work suggests that the nicotinic α5 subunit accelerates and strengthens the endogenous cholinergic response in prefrontal cortex and protects it against desensitization (22, 23). These nicotinic responses in pyramidal neurons are sensitive to aging and AD neuropathology (21, 71), but little is known about endogenous cholinergic modulation of chandelier neurons.

### Chandelier cells are potentially functionally affected by β-amyloid

Chandelier cells are a specialized subtype of PVALB+ interneurons. They differ anatomically from the PVALB+ basket cells by their large number of vertically oriented axonal cartridges, which specifically innervate the axon initial segments of pyramidal neurons (72,73). New evidence suggests chandelier cells regulate excitation dynamics of neuronal networks (74). Impairment of these neurons has been implicated in diseases involving pathological excitation in the cortex, such as epilepsy (75,76) and AD (3,4,68). Our RNASeq findings are in agreement with previous work showing that the inhibitory output of chandelier cells is sensitive to β-amyloid pathology (77) but unaffected by tau pathology (79). Chandelier cell axons near β-amyloid plaques have been found to show deformations, and pyramidal neurons proximal to plaques show loss of inhibitory input onto their axon initial segments (77). Our findings suggest that in people homozygous for A allele of the missense *CHRNA5* SNP rs16969968 (11% of the ROS/MAP participants), this vulnerability of chandelier cells to β-amyloid pathology may be exacerbated, possibly leading to cell death.

### Potential mechanisms for a neuroprotective effect of α5-containing nicotinic receptors

Our results suggest that polymorphisms affecting *CHRNA5* expression and function, may alter both the total β-amyloid levels in the brain, and alter the susceptibility of specific *CHRNA5*-expressing cell types, such as the chandelier cells, to β-amyloid-mediated toxicity. One possible explanation for these observations may be the lowered binding of β-amyloid to the α4β2α5 nicotinic receptors (14) expressed by these cells. This protection against β-amyloid binding and inhibition of the nicotinic response (13) could promote resilience of nicotinic signalling in the chandelier cells, potentially leading to improved cell survival (15) in AD pathology. Furthermore, since α4β2α5 nicotinic receptors support higher conductance of calcium ions into the cell (27,28), another putative neuroprotective mechanism of the α5 subunit may be through driving possible calcium-dependent neuroprotective pathways in neurons which express the α4β2α5 nicotinic receptors (79,80). Such calcium-regulated pathways may include *MME*, *SORL1*, *SPOCK1* or *PRNP*. These genes are specifically enriched in PVALB+ chandelier cells compared to PVALB+ non-chandelier cells (35) and have been previously suggested to alter β-amyloid production and clearance (53,56,57,81).

### Caveats and opportunities for additional investigation

While the ROS/MAP database offered an opportunity to assess the impact of *CHRNA5* expression and *CHRNA5*-related SNPs on AD pathology in a large sample, some caveats exist. *CHRNA5* expression has previously been shown to be important for animal performance in demanding attentional tasks (24,29), but a robust attention assessment of the ROS/MAP individuals was not part of the study design, potentially explaining the lack of any association between *CHRNA5* expression or polymorphisms and a cognitive readout. There is limited knowledge on the brain functions regulated by the *CHRNA5-*expressing chandelier cells. As a class, chandelier cells strongly regulate action potential generation at the axonal initial segment of pyramidal cells (72,73), potentially connecting them to seizure pathology (75,76). A previous study in rodents (69) has suggested a link between the rs16969968 SNP and certain features of psychosis, namely hypofrontality of schizophrenia patients (82). However, hypotheses relating to these potential links could not be assessed in our current study due to a lack of data on the diagnoses of the ROS/MAP participants for seizures or dementia-related psychosis. While the ROS/MAP dataset presented an opportunity to study the effects of rs16969968 on a background of an unusually-low smoking prevalence (83), future work would benefit from more robust assessment of smoking history.

Although most prevalent in the prefrontal cortex, the α4β2α5 receptor is not the only type of α5-containing nicotinic receptor. Another type of interest is the α3β4α5 receptor, which is expressed primarily in the habenula (28), and intriguingly has all its subunits within the same locus (83). Unfortunately, expression data for the relevant *CHRNA3* and *CHRNB4* genes were not included in the prefrontal bulk RNAseq dataset. β-amyloid pathology affects other types of nicotinic receptors besides the α4β2* subtype, including the widely-expressed low-affinity homomeric α7 receptor (11). However, since the activity and β-amyloid-sensitivity of the neuronal α4β2* receptor can be further modified by the inclusion of the auxiliary subunit α5 (14), we focus on the high-affinity nicotinic receptor as a potentially rewarding target of study in the context of altered nicotinic signalling in AD.

A limitation of the single-nucleus RNA sequencing data (36) was its relatively low number of individuals, limiting the robustness of comparing the effects of rs1979905 on *CHRNA5* expression across the different cell types, and preventing a similar examination of cell-type specific effects of rs16969968 on *CHRNA5* expression in the ROS/MAP dataset. This limitation should be considered when interpreting the findings of our study. Since cortical cell-type proportions were estimated using patterns of marker gene expression (36), the decreased chandelier cell proportions may instead reflect reductions of chandelier cell cartridges in AD (68). While the disruption of cortical E/I balance would remain similar, a different interpretation of our data would be that the rs16969968 A allele homozygous genotype exacerbates chandelier cartridge loss and results in lower expression of chandelier-cell-specific marker genes in individuals with elevated β-amyloid. Furthermore, while SNP exploration provides novel insight into the relationship between *CHRNA5* and neuropathology in aging, this work is correlational. Gene expression does not necessarily denote protein levels in AD brains (85) and thus differences in nicotinic receptor gene expression may not fully predict receptor levels or binding (86). Work in model systems and larger snRNAseq datasets will be necessary to test specific hypotheses raised in this work.

Finally, our preliminary analysis using the ADNI dataset did not show a significant link between *CHRNA5* SNPs and amyloid pathology, like that which we found in the larger ROS/MAP dataset. However, a comparison between these findings is complicated by the major differences between the amyloid level measurements methods utilized by the two studies. While our analyses in ROS/MAP used the clinical gold standard for the assessment of neuropathology in the brain, namely *post-mortem* immunohistochemical analysis, the amyloid measurements in ADNI are based on positron emission topography (PET) and spinal tap sampling of cerebrospinal fluid (CSF). Future studies, with a more highly powered ADNI dataset and an increased understanding of premortem and postmortem amyloid assessments, could further build on this work.

### Summary and implications

A growing body of work suggests that cortical excitability is perturbed early in Alzheimer’s disease through impairment of inhibitory interneurons (63,87). *CHRNA5* is positioned to modulate the overall excitability of the prefrontal cortex in two ways: through excitation of a population of deep layer cortical pyramidal neurons (22,23,88,89) that send projections throughout prefrontal cortex (23,90) and, as ascertained from our findings in this study, through excitation of a specific subset of cortical interneurons, the chandelier cells. Our findings suggest that *CHRNA5* is involved in Alzheimer’s disease neuropathology. The A allele of the *CHRNA5* regulatory-SNP rs1979905 predicts higher expression of *CHRNA5* and reduced β-amyloid load in the brain. In parallel, the A allele of the missense SNP, rs16969968, is associated with fewer chandelier cells in individuals with high β-amyloid levels, suggesting that differences in the trafficking of *CHRNA5* alter cellular resiliency to β-amyloid pathology. This combination suggests neuroprotective roles of *CHRNA5* in β-amyloid pathology and makes *CHRNA5* a target for therapies aiming to improve neuron survival in Alzheimer’s disease.

## Acknowledgements

This work was supported by the Canadian Institutes of Health Research (CIHR; PJT-153101, EKL; MOP89825, EKL; NGN-171423, ST; PJT-428404, DF), Krembil Foundation (DF, ST), Koerner Family Foundation (DF), CAMH Discovery Fund (DF, ST), Kavli Foundation (ST), McLaughlin Foundation (ST), Natural Sciences and Engineering Research Council of Canada (RGPIN-2020-05834 and DGECR-2020-00048; ST). JM is supported by the Canadian Research Chairs program, as a Tier 1 Chair. ROS/MAP is supported by P30AG10161, P30AG72975, R01AG15819, R01AG17917. U01AG46152, U01AG61356. ROS/MAP resources can be requested at https://www.radc.rush.edu

