## Supplemental Materials for "*CHRNA5* links chandelier cells to severity of amyloid pathology in aging and Alzheimer’s Disease"

- **Supplemental Methods**

- **Supplemental Figure S1**
- **Supplemental Figure S2**
- **Supplemental Figure S3**

- **Supplemental Table S1**
- **Supplemental Table S2**
- **Supplemental Table S3**
- **Supplemental Table S4**

- **Supplement References**

### Supplemental Methods

#### Genotype data preparation and imputation

ROS/MAP participants were genotyped in two batches (Synapse ID: syn3219045): the first batch (n=1709) using the Affymetrix GeneChip 6.0 (Affymetrix, Inc, Santa Clara, CA, USA) at the Broad Institute's Center for Genotyping or the Translational Genomics Research Institute, and the second batch (n=382) using the Illumina HumanOmniExpress (Illumina, Inc, San Diego, CA, USA) at the Children's Hospital of Philadelphia. Both batches underwent the same initial quality control (QC) analysis, as described previously (1). Only individuals with self-reported European ancestry were genotyped to minimize population heterogeneity. Sample-level quality control assessment included exclusion of samples with genotype success rate <95%, discordance between inferred and reported gender, and excess heterozygosity. Imputation of genotype data was performed as described previously (2). Each batch was imputed separately using the TOPMed Imputation Server (TOPMed reference r2), including Eagle (v2.4) for allelic phasing and Minimac4 (v1.5.7) for imputation. Prior to submission for imputation, genotypes were pre-processed using the TOPMed Imputation Server-recommended data preparation pipeline available here: https://topmedimpute.readthedocs.io/en/latest/prepare-your-data.html. Imputed output data from the TOPMed server for each batch were filtered for imputation quality (removing SNPs with r<0.8) before merging and mapping to rsIDs (dbSNP build 155). This resulted in a final high-quality dataset of 9,329,439 bi-allelic autosomal SNPs from 2061 individuals. Minor allele frequencies of candidate SNPs were calculated using the getmaf function of the BinaryDosage (1.0.0) R package.

#### Fine population structure

Principal component analysis (PCA) was performed on genotyped SNPs using PLINK2 (3) to derive principal components representing fine population structure. The first 10 principal components were used as covariates in all linear regression models that included SNP or gene expression data.

#### GWAS summary statistics of *CHRNA5* SNP association with Alzheimer's disease

Summary statistics were downloaded from the NHGRI-EBI GWAS Catalog (4) on 31/10/2023 for study GCST90027158 (5). Original GWAS p-values were adjusted using the Bonferroni correction method (6) using the p.adjust function in R. Detailed methods of the GWAS generation are available in Bellenguez et al. 2022 (5).

#### Neuropathology and cognitive scores

A detailed description of the neuropathology and cognitive variables in ROSMAP is included on the RADC Research Resource Sharing Hub. The ROS/MAP neuropathology data used in our study included measurements of average β-amyloid protein load and the density of paired helical filament tau tangles across 8 brain regions (hippocampus, entorhinal cortex, midfrontal cortex, inferior temporal cortex, angular gyrus, calcarine cortex, anterior cingulate cortex, superior frontal cortex). Amyloid beta protein was identified by molecularly specific immunohistochemistry and quantified by image analysis as described previously (7) as percent area of cortex occupied by amyloid beta. Neuronal neurofibrillary tangles were identified by molecularly specific immunohistochemistry (antibodies to abnormally phosphorylated Tau protein, AT8) as described previously (7). Cortical density (tangle counts per mm2) was determined using systematic sampling. As recommended by the ROS/MAP study authors, the square-root of the averages of β-amyloid load and tangle density was used in all analyses in our study. Post-mortem neuropathology data was available from 1694 individuals.

The global cognition score was calculated at each visit as the average of 19 tests assessing the individuals' episodic, working, and semantic memory, and perceptual orientation and speed. The raw score of each test were converted to Z scores and averaged to yield a global cognition score. The z-scores have a mean of 0 and standard deviation of 1. A negative z-score means that someone has a global cognition score lower than the cohort average at baseline. We used only the global cognition score recorded at the subject's last recorded session before death for our analysis. Global cognition scores at the last visit before death were available from 1853 individuals.

**Smoking status**

The smoking status of the ROS/MAP participants was determined at the baseline interview by participants' answers to the following questions: 1. "Do you smoke cigarettes now?"; 2. "Did you ever smoke cigarettes regularly". Question 1 was used to determine current smokers, and question 2 identified former smokers. Participants who answered "no" to both questions were categorized as "never smoked".

**Tissue preparation and bulk tissue RNA sequencing**

Details on the preparation and handling of ROS/MAP post-mortem samples have been previously published (8). Samples were extracted using Qiagen's miRNeasy mini kit (cat. no. 217004) and the RNase free DNase Set (cat. no. 79254) and quantified by Nanodrop and quality was evaluated by Agilent Bioanalyzer. RNA sequencing on DLPFC tissue was carried out in 13 batches within three distinct library preparation and sequencing pipelines. Sequencing was carried out using the Illumina HiSeq (pipeline #1: 50M 101bp paired end reads) and NovaSeq6000 (pipeline #2: 30M 100bp paired end; pipeline #3: 40-50M 150bp paired end 121 reads). Full details on RNA extraction and sequencing are available on the Synapse AMP-AD 122 Knowledge Portal (syn3219045) and in Consens et al., 2022.

**Processing of bulk DLPFC tissue RNAseq dataset**

DLPFC RNAseq data processing, including gene expression residuals generation was performed as previously described (9). In brief: All 13 DLPFC batches (ntotal = 1110) were processed using the same pipeline: 1) fastq file quality control was performed using FastQC v0.11.5 (default parameters), 2) STAR v2.5.3a was used to align reads (GRCh38.91 reference), 3) RSEM v1.2.31 was used to quantify expression from aligned BAM files, and 4) multiqc v1.5 was used to aggregate quality metrics from fastqc and Picard tools v2.17.4, 5) quality reports were examined for each batch and exclusion of samples was initially carried out according to manual identification of outlying samples primarily considering low numbers of aligned reads, excess GC coverage bias, high percentage of read duplicates, and abnormal distribution of read assignments across genomic annotations. Expression of the *XIST* gene was evaluated at this stage to exclude subjects with contradictory reported sex, 13 subjects were identified (new n total=1097).

***DLPFC RNAseq data quality control***

Full details of QC are available in Felsky et al. 2022. In brief: expected counts, calculated by RSEM, were aggregated, and used as input to limma (v3.48.3) voom in R (v4.1.1). Genes with insufficient expression (median count value was less than 15 across the combined sample) were removed. Naïve multidimensional scaling (MDS) analysis was then performed on the top 5000 most variable genes (using limma "plotMDS") to identify subjects with outlying expression patterns. Outliers were defined as those with values of either of the first two latent dimensions exceeding ±4 times the interquartile range (IQR) of their within-batch median value. Following this step, 1091 subjects remained for the DLPFC data. These QC data were then trimmed mean of M-values (TMM) normalized.

***Final QC and generation of gene expression residuals***

To ensure the robustness of included data, linear effects of identified technical co-variates, age at death and post-mortem interval (PMI), and biological sex were removed using the "lmfit" (specifying robust Huber regression (10), while batch effects were removed using the removeBatchEffect function in limma (11), and expression values residualized. Individuals were then hierarchically clustered (agglomerative) and the resulting dendrograms (Supplementary figure 9 of Felsky et al. 2022) were manually inspected to identify any additional subject outliers escaping recognition by two-dimensional MDS (18 identified). This resulted in a high-quality dataset of DLPFC bulk tissue ($n_{genes}$=17 465). The calculated gene expression residuals were used in all analyses assessing gene expression with other variables in our study.

**Cohort characteristics for the DLPFC single-nucleus RNA sequencing data**

The DLPFC snRNAseq data published by Cain et al. 2020 was generated from 24 ROSMAP participants (12). Since lack of matching with genotyping data excluded 2 participants, we used the snRNAseq data from the remaining 22 in our analysis. The participants included were balanced for sex and disease status. 50% of participants were female and the diagnosis for this group was 50% NCI and 50% AD as determined by final clinical consensus. The average age at death was 87.96 ± 5.64 years and the average PMI was 5.03 ± 1.93 hr. Average brain neuropathology levels were amyloid 1.23 ± 1.09 and tangles 2.093+/-1.221.

**Single-nucleus RNA sequencing and pre-processing**

Details on the nucleus isolation and single nucleus RNA sequencing from the dorsolateral prefrontal cortex of 24 ROS/MAP individuals performed by Cain et al. 2020 are available on the Synapse AMP-AD Knowledge Portal (ID: syn16780177) and in the Cain et al. 2020 preprint. In brief, nuclei from an individual sample were loaded on a single channel on the 10x Genomics Chromium chip. Library construction was then performed as per the Chromium Single Cell 3' Reagent Kit v2 protocol (CG00052) provided by the manufacturer. The described protocol produces Illumina-ready sequencing libraries. All sequencing was performed on an Illumina HiSeq4000 machine. Generation of count files from the raw sequencing data was performed using the Cellranger software package (version 2.1.1, chemistry V2, 10X Genomics, for chemistry Single Cell 3'). For every nucleus, the number of genes for which at least one read was mapped was quantified, all nuclei with fewer than 400 detected genes were excluded.

**Estimations of cell type proportions for bulk DLPFC RNAseq data**

Estimates of cell-type proportions from bulk DLPFC RNAseq data from 640 ROS/MAP participants was performed and described by Cain et al. 2020. To achieve this the authors developed a regression-based consensus model (CelMod) which leveraged the matched snRNAseq and bulk RNAseq data from the same 24 ROSMAP participants. The authors first trained the model by 1) In the snRNAseq data,

performing a linear regression on each gene separately for each cell cluster of interest, using its expression as the dependent variable and the proportion of that cluster in each snRNA-Seq sample in the training set as the independent variable. 2) In the matched bulk RNAseq data, for each gene using the regression model to calculate the predicted proportion of each cell type, normalizing their sum to 1. 3) Ranking the genes by the 90th percentile of the absolute value of the error between predicted and training proportions, for each cell type. 4) Selecting the number of top-ranked genes to use for deconvolving a new bulk RNAseq sample. The authors then used these top-ranked genes to estimate the cell type proportions in the larger (non-snRNAseq-matched) bulk RNAseq dataset.
The authors ran this algorithm in an iterative manner starting from broad cell classes (glutamatergic neurons, GABAergic neurons, astrocytes, oligodendrocytes, OPCs, microglia, endothelial cells, and pericytes) and then for the subtypes within each of these higher-level classes. For the broad cell classes, the proportions were based on the total nuclei per sample. For the subtypes, the proportions were normalized to the total nuclei from the broad class of interest - for example, for the GABAergic neuron subtypes, the proportions for the training (and thus the prediction) were normalized to the total number of GABAergic neuron nuclei per sample.

**ADNI database methods and analysis**

The Alzheimer's Disease Neuroimaging Initiative (ADNI)(13) is an ongoing longitudinal study composed of four phases (ADNI-1, ADNI-GO, ADNI-2, and ADNI-3) and enrolling groups of participants with clinical Alzheimer's disease, early and late mild cognitive impairment, significant memory concern, or no cognitive impairment. Full study details are available at https://adni.loni.usc.edu/ . All Participants provided informed consent. Clinical, imaging, fluid biomarker, and genetic data are collected at baseline and at 6 and 12-month follow-ups. Genotype data for participants were obtained at every phase: $n_{ADNI-1}$=757 genotyped using the Illumina Human610-Quad BeadChip, $n_{ADNI-GO/2}$=432 genotyped using the Illumina HumanOmniExpress BeadChip, and $n_{ADNI-3}$=327 genotyped using the Illumina Global Screening Array v2. Standard quality control was applied to the genotype data before imputation was performed on the TOPMed Imputation Server, resulting in 8,028,924 high-quality variants in n=1,569 participants. Principal component analysis (PCA) was performed on genotyped SNPs using PLINK2 (3).
The ADNI dataset includes measures of premortem levels of brain β-amyloid levels (n=577) measured using PET imaging following florbetapir or florbetaben injection. Details on the PET scan procedure and quantification are available on the ADNI website. ADNI also includes measurements of β-amyloid in the CSF collected through lumbar puncture (n=637) and measured using ELISA.
Associations between the *CHRNA5* SNPs (in both sample-major additive and dominant genotype coding) and brain or CSF β-amyloid levels were assessed using linear regression models, separately for rs1977905 and rs16969968, co-varying for age, sex, and the first ten genomic principal components.

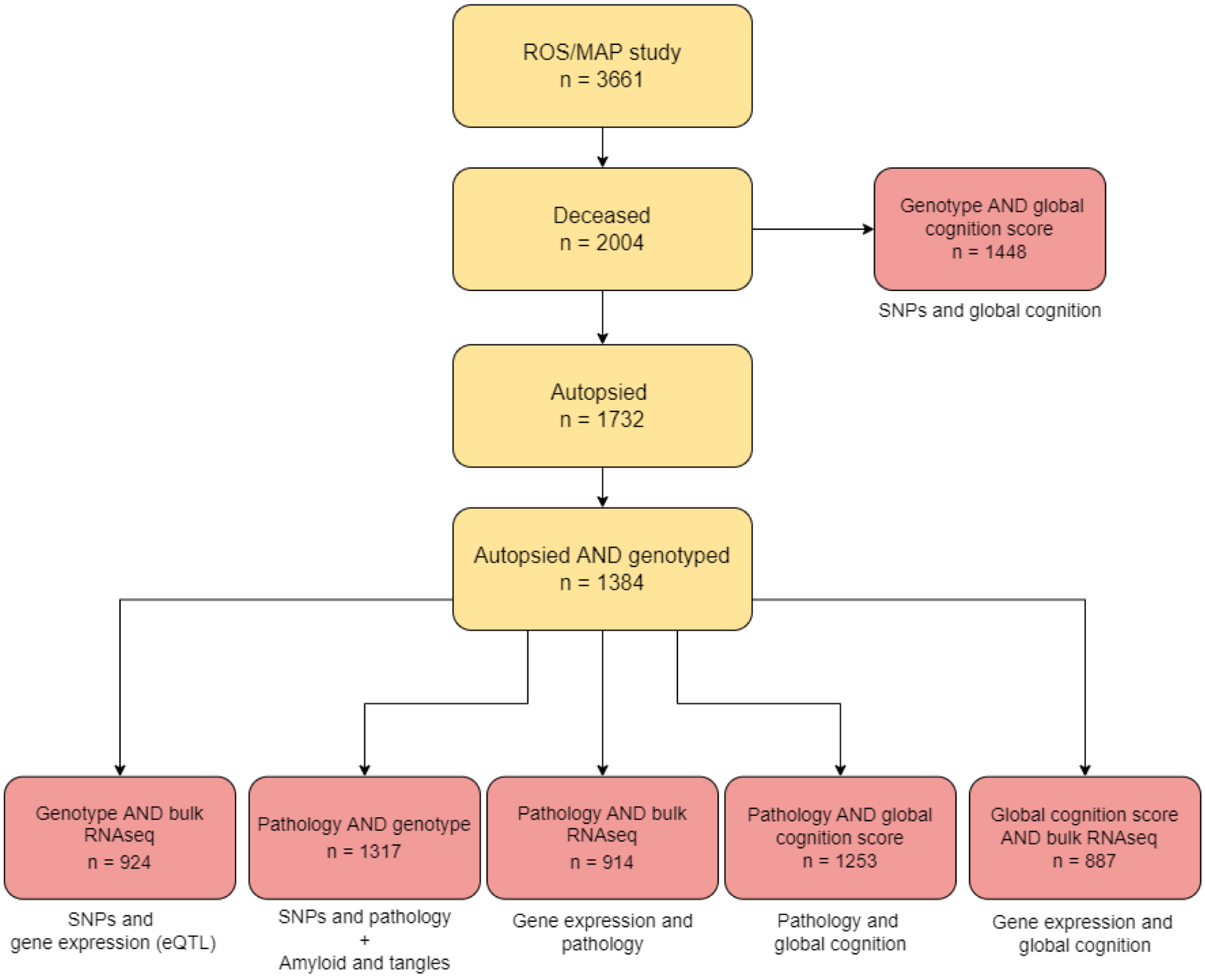


**Supplemental Figure S1. A flow chart describing the ROSMAP participants we used in the different linear models in our analysis.** Boxes indicate populations of ROSMAP participants with the number remaining in each exclusion step (*n*). The text under individual boxes describes the associations which were tested using the specific ROSMAP population in the box above.

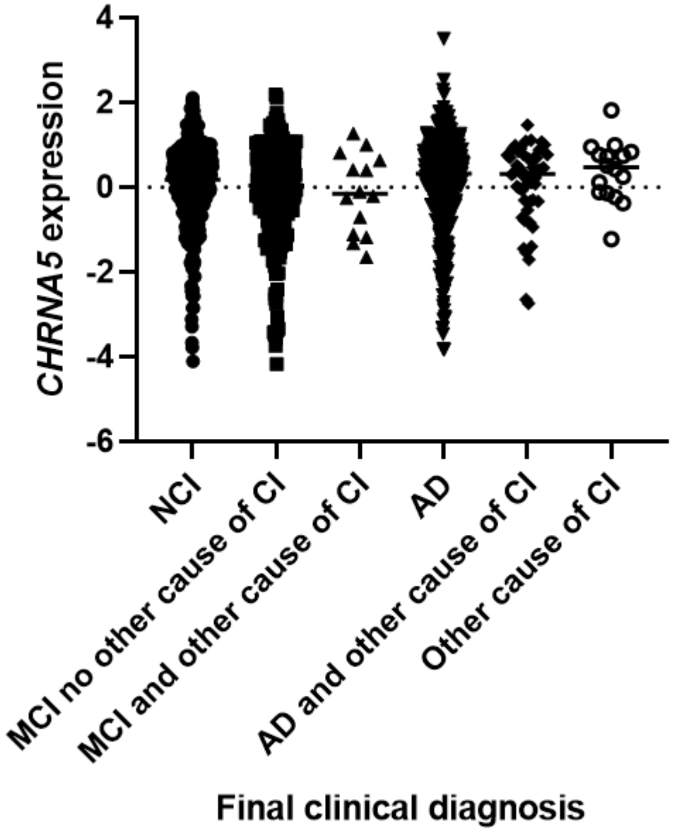


<u>Supplemental Figure S2</u>. **A distribution plot of *CHRNA5* expression across different disease states as determined by the final clinical diagnosis.** No significant difference was found in *CHRNA5* gene expression in the bulk DLPFC between the different disease states as determined by final consensus clinical diagnosis (One-way ANOVA with Tukey's *post-hoc t*-test: F = 1.693, $p$ = 0.134). No cognitive impairment (NCI), mild cognitive impairment (MCI), cognitive impairment (CI), Alzheimer's Disease (AD).

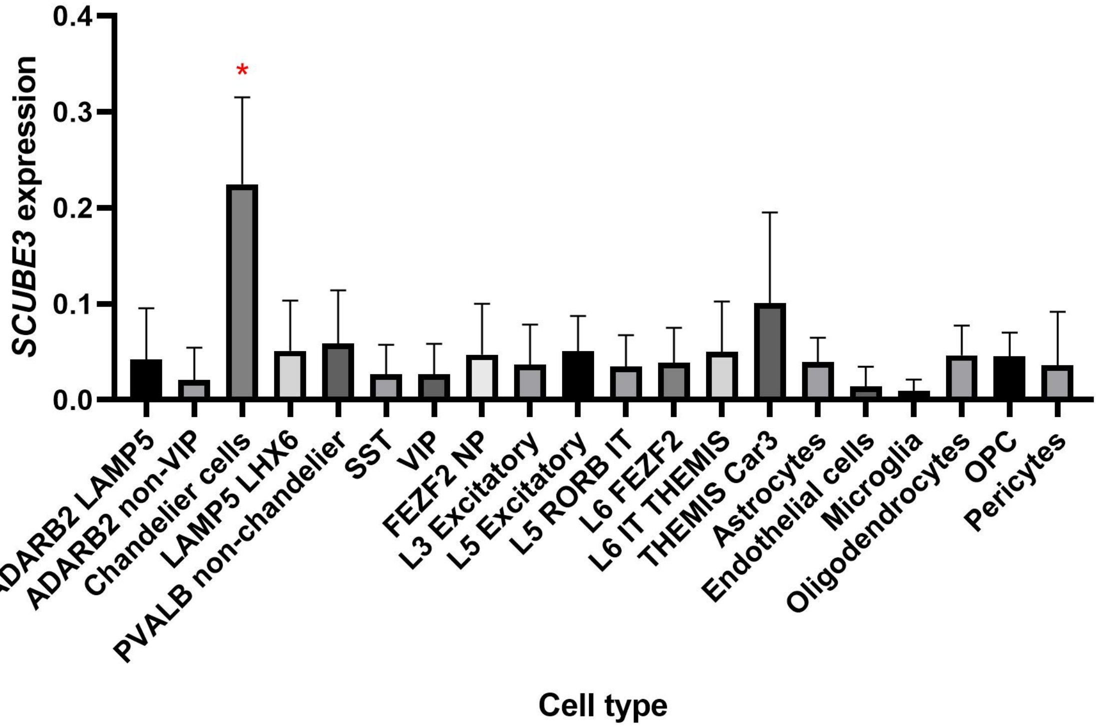


**Supplemental Figure S3. *SCUBE3* expression is elevated in chandelier cells.** Expression of *SCUBE3* averaged per cell type per individual, original gene count values were normalized for each cell by total expression. F-test significance of ANOVA shown on graph, with red asterisk denoting post-hoc tests demonstrating *SCUBE3* expression is stronger in chandelier cells compared to all other cell types ($p < 0.0001$ for all comparisons).

**Supplemental Table S1**. Number of cells per cell subtype cluster in the genotype-matched snRNAseq data from Cain et al 2020.

| **Cell type** | **Number of cells** |
|---|---|
| ADARB2 LAMP5 | 1733 |
| ADARB2 non-VIP | 1822 |
| Chandelier cells | 1117 |
| LAMP5 LHX6 | 853 |
| PVALB non-chandelier cells | 6476 |
| SST | 4302 |
| VIP | 5780 |
| FEZF2 NP | 1460 |
| L3 Excitatory | 24232 |
| L5 Excitatory | 8942 |
| L5 RORB IT | 9943 |
| L6 FEZF2 | 2812 |
| L6 IT THEMIS | 2729 |
| THEMIS Car3 | 892 |
| Astrocytes | 27461 |
| Endothelial cells | 1857 |
| Microglia | 3829 |
| Oligodendrocytes | 32781 |
| OPC | 6835 |
| Pericytes | 1078 |

**Supplemental table S2**: Results of linear regression of rs1979905 A genotype and *CHRNA5* expression in each cell type (in the snRNAseq dataset). Significant results ($p < 0.05$) are italicised. Pericytes showed no *CHRNA5* expression.

| **Cell type** | **Association with rs1979905 A: t value (p value)** |
|---|---|
| ADARB2 LAMP5 | -0.117 (0.909) |
| ADARB2 non-VIP | 0.506 (0.620) |
| Chandelier cells | 0.554 (0.587) |
| LAMP5 LHX6 | 0.804 (0.433) |
| PVALB non-chandelier | 0.683 (0.504) |
| SST | 0.982 (0.339) |
| VIP | -0.092 (0.927) |
| FEZF2 NP | 0.535 (0.6) |
| L3 Excitatory | 0.322 (0.751) |
| L5 Excitatory | -0.112 (0.913) |
| L5 RORB IT | *2.460 (0.025)* |
| L6 FEZF2 | 0.762 (0.456) |
| L6 IT THEMIS | *2.402 (0.028)* |
| THEMIS Car3 | 0.048 (0.963) |
| Astrocytes | 1.229 (0.236) |
| Endothelial cells | 0.224 (0.825) |
| Microglia | 0.812 (0.428) |
| Oligodendrocytes | *2.416 (0.027)* |
| OPC | *2.351 (0.031)* |
| Pericytes | NA |

**Supplemental Table S3.** Summary of our findings on the *CHRNA5* SNPs of interest (rs1979905 and rs16969968).

| **Findings** | **Rs1979905_A (regulatory)** | **Rs16969968_A (coding)** |
|---|---|---|
| eQTL | Significant association with *CHRNA5* expression in the prefrontal cortex in bulk and single-cell RNAseq. No effect on *CHRNA5* expression in chandelier cells (show highest expression of *CHRNA5*). | No association with *CHRNA5* expression once adjusted for rs1979905 genotype. |
| Neuropathology | Significantly associated with lower levels of cortical amyloid levels. | Significantly affects association between cortical amyloid levels and proportions of chandelier cells in the prefrontal cortex |

**Supplemental Table S4.** Summary GWAS statistics of the association of rs1979905 and rs16969968 with Alzheimer's disease from Bellenguez et al. 2022. $P_{Bonf}$ is original regression p value adjusted using the Bonferroni correction method. Beta-coefficient and standard error (SE) are reported.

| Chromosome | Position | SNP | Effect allele | Non-Effect allele | Beta | SE | P | $P_{Bonf}$ |
|---|---|---|---|---|---|---|---|---|
| 15 | 78842374 | rs1979905 | A | C | 0.02 | 0.008 | 0.015 | 1 |
| 15 | 78882925 | rs16969968 | A | G | -0.023 | 0.009 | 0.007 | 1 |
